# Non-canonical secondary structures arising from non-B-DNA motifs are determinants of mutagenesis

**DOI:** 10.1101/146621

**Authors:** Ilias Georgakopoulos-Soares, Sandro Morganella, Naman Jain, Martin Hemberg, Serena Nik-Zainal

**Author notes:** Corresponding authors: Martin Hemberg & Serena Nik-Zainal.

## Abstract

Somatic mutations show variation in density across cancer genomes. Previous studies have shown that chromatin organization and replication time domains are correlated with and thus predictive of this variation ^1,2,3,4,5^. Here, we analyse 1,809 whole-genome sequences from nine cancer types ^6,7,8^ to show that a subset of repetitive DNA sequences called non-B motifs that predict non-canonical secondary structure formation ^9,10,11,12^ can independently account for variation in mutation density. However, combined with epigenetic factors and replication timing, the variance explained can be improved to 43-76%. Intriguingly, ~2-fold mutation enrichment is observed directly within non-B motifs, is focused on exposed structural components, and is dependent on physical properties that are optimal for secondary structure formation. Therefore, there is mounting evidence that secondary structures arising from non-B motifs are not simply associated with increased mutation density, they are possibly causally implicated. Our results suggest that they are determinants of mutagenesis and increase the likelihood of recurrent mutations in the genome ^13,6^. This analysis calls for caution in the interpretation of recurrent mutations and highlights the importance of taking non-B motifs, that can simply be inferred from the reference sequence, into consideration in background models of mutability henceforth.

## Main Text

The canonical right-handed DNA double-helical structure, known as B-DNA, has been recognized since 1953. Although B-DNA is the predominant configuration inside the cell, more than 20 non-canonical secondary structures have been reported^9^. These alternative structures include triple-helices, hairpins, cruciforms and slipped structures, and they are more likely to form at particular repetitive sequences such as mirror repeats, inverted repeats, direct repeats and short tandem repeats ^10^. Non-canonical secondary structures are associated with increased mutability according to *in vitro* studies of prokaryotic ^14,15^ and eukaryotic cells ^16,17,18,19,20,21,22,23,24,25,26^. Here, we methodically explore the relationship between secondary structures and somatic mutability, focusing on seven common types of sequence motifs prone to forming non-canonical secondary structures, hereafter referred to as non-B DNA motifs for brevity: direct repeats (DR), G-quadruplexes (G4), inverted repeats (IR), mirror repeats (MR), H-DNA, short tandem repeats (STR) and Z-DNA (Fig. 1a-f, definitions of each of these can be found in Methods).

**Figure 1:**
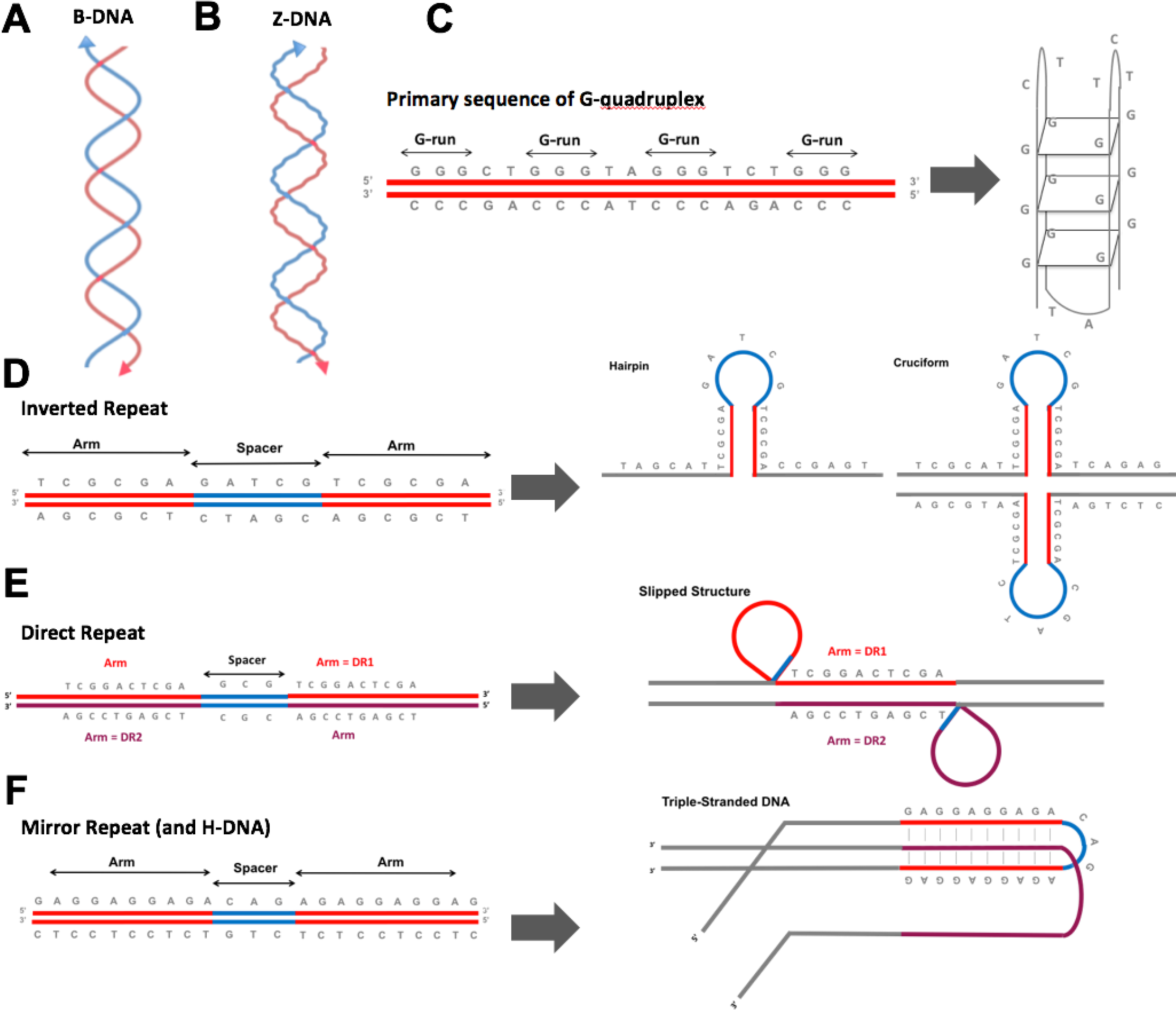
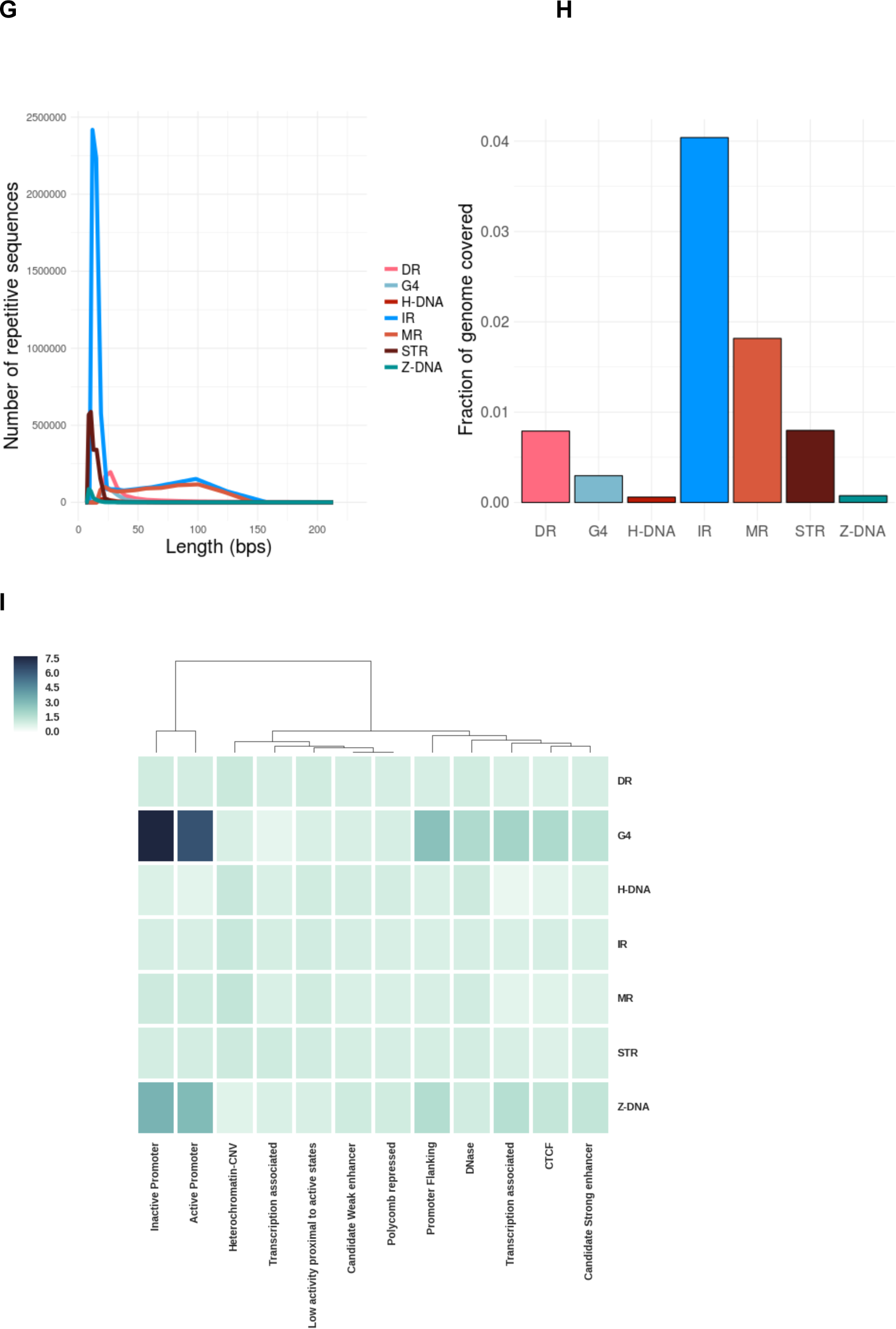
**Non-canonical secondary structures arising from non-B DNA motifs in the human genome** *(a)* Normal configuration of human DNA. (b) Left-handed helical structure caused by Z-DNA *(c-f)* Schematic representations of the primary sequence of various non-B motifs and their corresponding predicted secondary structures. (g) Distribution of lengths of non-B-DNA motifs. *(h)* Fraction of the human reference genome (hg19) covered by different non-B-DNA motifs. *(i)* Enrichment of occurrences of non-B DNA motifs associated with various chromatin states (Methods for calculation).

We systematically explored each of the seven non-B DNA motifs in the human reference sequence (Methods)^11^. Most motifs are <50 bps (Fig. 1g), and each category encompasses 0.07% to 4% of the human genome (Fig. 1h) which may seem small fractionally, but absolute numbers of motifs are substantial (range 69,154-6,006,266). Non-B motifs show non-uniform distributions across the genome reflected by their variable enrichments at different chromatin-associated regions (Fig. 1i): G4 and Z-DNA are strongly enriched at GC-rich promoter regions; DR, H-DNA and MR are modestly enriched in low complexity repetitive sequences (e.g. heterochromatin); and IR and STR are more uniformly distributed between gene-rich through to gene-poor regions. Although some motifs are correlated with each other, there is limited overlap between distinct types of non-B motifs (Fig. S1 a-b).

Genomic features such as histone epigenetic marks and replication time domains have been shown to be predictive of the variation in distribution of somatic mutations ^1,2^. We thus explored whether non-B motifs also had an impact on somatic mutagenesis. We used mutation catalogues derived from 560 whole-genome sequenced (WGS) breast cancers^6^. The genome was binned, and mutations, non-B motifs, histone modifications and replication time domains were counted for each bin (methods, Fig. S2, Fig. S3). Consistent with previous reports ^1,2,27,3,4^, we find that genomic features linked to epigenetic modifications such as heterochromatin (H3K9me3, r=0.31) and late replicating domains (r=0.59) are associated with increased mutational density while open chromatin (DNASE r= −0.31), active cis-regulatory elements (H3K27ac, r= −0.52) and transcribed regions (H3K36me3, r= −0.57) are negatively associated with mutational density (Fig. 2a, Fig. S4). Surprisingly, crude correlations for selected non-B DNA motifs, particularly IR (r=0.28), STR (r= −0.33), G4 (r= −0.38), MR (r= 0.20) and Z-DNA (r= −0.19) (Fig. 2a, Fig. S4) are seen. Partial correlation analysis reveals the association between somatic mutations and non-B motifs remains even after controlling for epigenetic marks and replication timing (Fig. S5), raising the possibility that non-B motifs are independent factors that could contribute to mutability ^28,29,23,30^.

**Figure 2:**
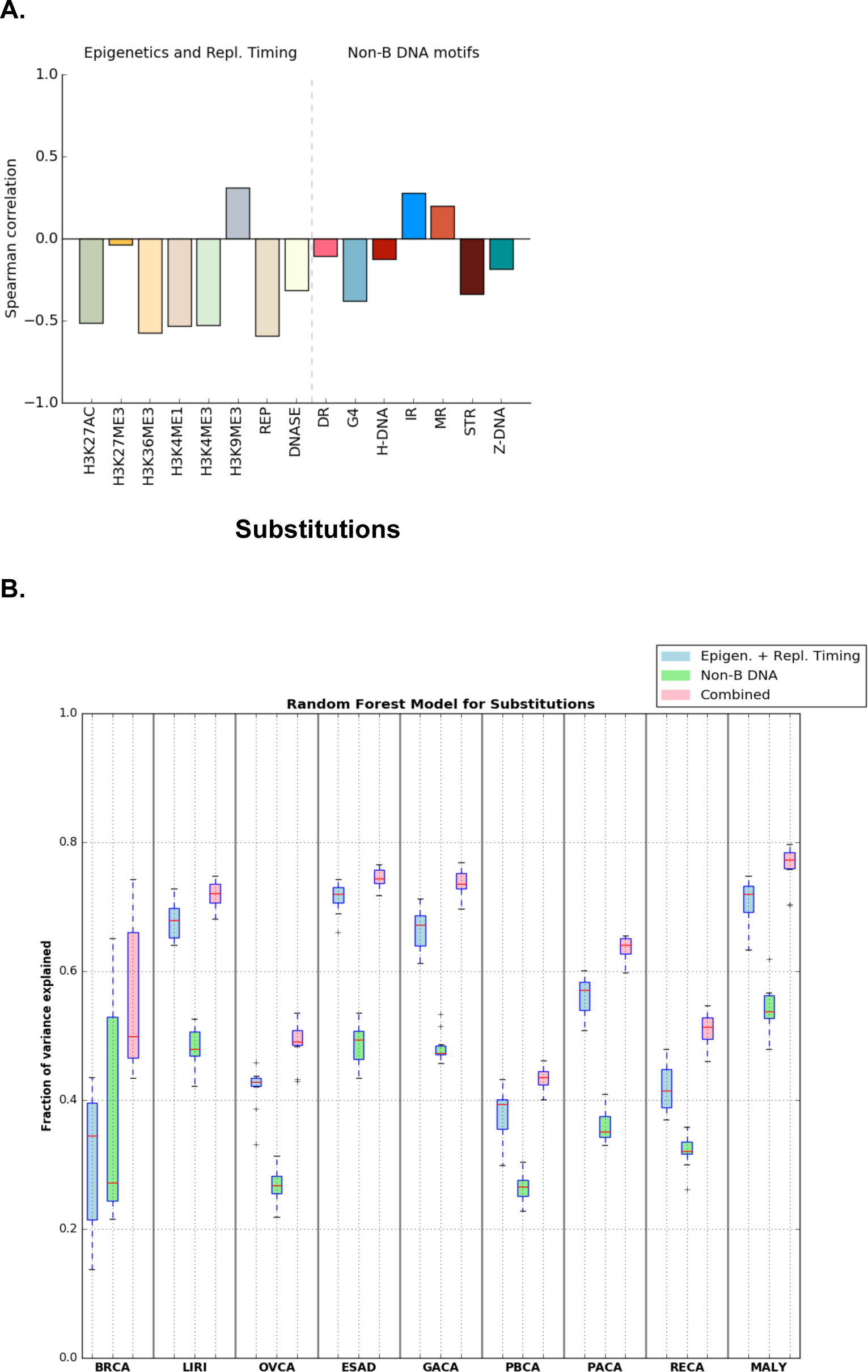
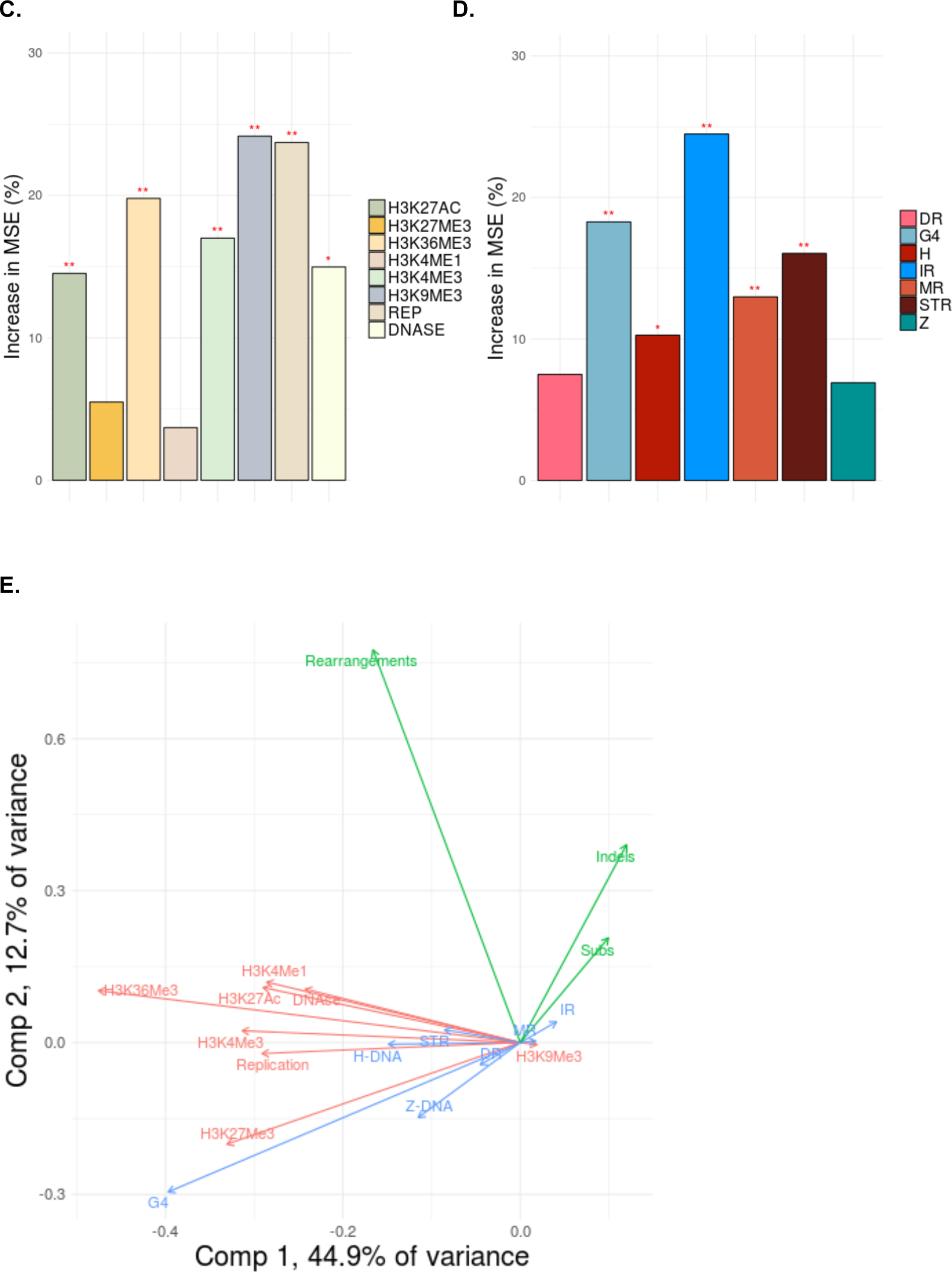
**Non-B-DNA motifs predict somatic mutability in human cancers *(a)*** Correlations between the number of non-B-DNA motifs, and epigenetic features and replication timing, with the number of substitutions (Spearman rank). Please note interpretation is directional e.g. A positive correlation with replication time would indicate increased mutability with early replication time domains, while a negative correlation denotes increased mutability in late replication time domains. *(b)* Fraction of variance explained for predicting the number of mutations in 500 kb bins with random forest regression using non B-DNA motifs and epigenetic features/replication timing as predictors for multiple tumor types (BRCA = breast cancer, LIRI=liver cancer, OVCA = ovarian cancer, ESAD = esophageal adenocarcinoma, GACA = gastric cancer, PBCA = pedriatic brain cancer, PACA = pancreatic cancer, RECA = renal cell carcinoma, MALY= malignant lymphoma). Error bars represent standard error from 10-fold cross-validation. *(c, d)* Importance of the different predictors for the random forest regression. The y-axis shows the increase in mean square error (MSE) when the variable is excluded. Bars with * have an FDR<.05 and ** have FDR<.01 as determined by a permutation test. *(e)* PCA Analysis. The first two principal components separate mutations (green), non-B DNA motifs (blue) and epigenetics and replication timing domains (red).

To explore this in more depth we assessed the predictability of mutation density given the number of non-B motifs (as well as epigenetic features and replication time domains) by constructing models using linear regression and random forest regression (Fig. 2b, Fig. S6). First, our analysis recapitulates previous studies showing that random forest regression is more accurate than linear regression for base substitutions and also identifies H3K9me3 and replication timing as the most informative features for predicting mutability ^1,2^ (Fig. 2c, Fig. S7). Second, we find that IRs and G4s are relatively strong predictors of mutability although other non-B motifs including MR, H-DNA, STR and Z-DNA contribute predictive power (Fig. 2d, Fig. S7). Third, although non-B motifs alone can explain 37% of observed variance in mutation density for base substitutions in breast cancer, regression models incorporating both epigenetic, replication time and non-B motifs substantially improve the variance explained to 52%, performing better than either model separately (Fig. 2b). The enhanced model predictions featuring combined data is unsurprising in light of a principal component analysis biplot: Non-B motifs and epigenetic features are separated by the second component (Fig. 2e) suggesting that they contribute towards predicted mutability in different ways, reinforcing the partial correlation analyses described earlier. Since non-B motifs can be computed from the reference genome alone, our results suggest a straightforward and cost-effective way of improving mutability predictions.

To validate our predictive model, we employed it across WGS cancer datasets from eight other tissue types including liver, ovarian, esophageal, gastric, pancreatic, renal cell carcinoma and pediatric brain cancers and malignant lymphoma ^31,32,33,7,8^. The predictive accuracy of the regression model varied by cancer type, with between 43-76% of the variance explained (Fig. 2b-d, Fig. S6-S7). Consistently across all tumour types, non-B motifs made a smaller independent contribution towards predicting mutability, but in combination with epigenetic factors/replication timing, improved predictive ability overall. Regression analyses were performed for other mutation classes - indels and rearrangements - and predictive ability of the model similarly improved when the factors were combined (Fig. S8). The model performed better for indels than for rearrangements, although the number of rearrangements is much lower than substitutions and indels (by orders of magnitude), hence we cannot exclude the possibility that model performance is limited by sample size. Our findings bring together and reinforce previous observations of indel enrichment at disparate non-B motifs in experimental systems (e.g. IRs ^34,35,36,37^; DRs ^38,39^; and G4s ^40,35,41^.

We thus conclude that primary sequence features, as represented by non-B DNA motifs, are collectively informative for predicting local mutability across many tissue types, predominantly of substitutions and indels. Is it the physical presence of a non-canonical secondary structure that mechanistically drives the increased likelihood of mutagenesis? The evidence in favour of this possibility is described below.

First, we find that somatic mutations are not simply increased in the vicinity of non-B motifs, they are elevated within non-B motifs themselves (Fig. 3a-b). H-DNA, STR and Z-DNA motifs were most enriched for substitutions 1.7-fold, 1.6-fold and 1.7-fold respectively, while other motifs showed more modest enrichment: G4 (1.2-fold), IR (1.1-fold), DR (1.1-fold) and MR (1.1-fold) when compared to their immediate surrounding sequence (i.e. corrects for genomic GC variation). There is more striking enrichment of indels in general: Z-DNA (10.7-fold), H-DNA (6fold), STR (5.8-fold), MR (2.5-fold), DR (2.3-fold) and G4 (1.5-fold), a finding that is not surprising given that most indels in human cancer occur at repeat tracts which are present at a higher frequency particularly at Z-DNA, H-DNA and STRs. For rearrangements, enrichment was observed within IRs in breast cancer (1.2-fold) (Fig. 3a) reinforcing observations in yeast and mammalian *in vitro* studies^17^. Enrichment of mutagenesis within non-B motifs is remarkably consistent across all tumor types for some motifs (e.g. Z-DNA, STR, G4, H-DNA) (Fig. S9). Essentially, we find that there is an excess of mutability not just associated with non-B motifs, but directly within them (Fig. 3a, Fig. S9).

**Figure 3:**
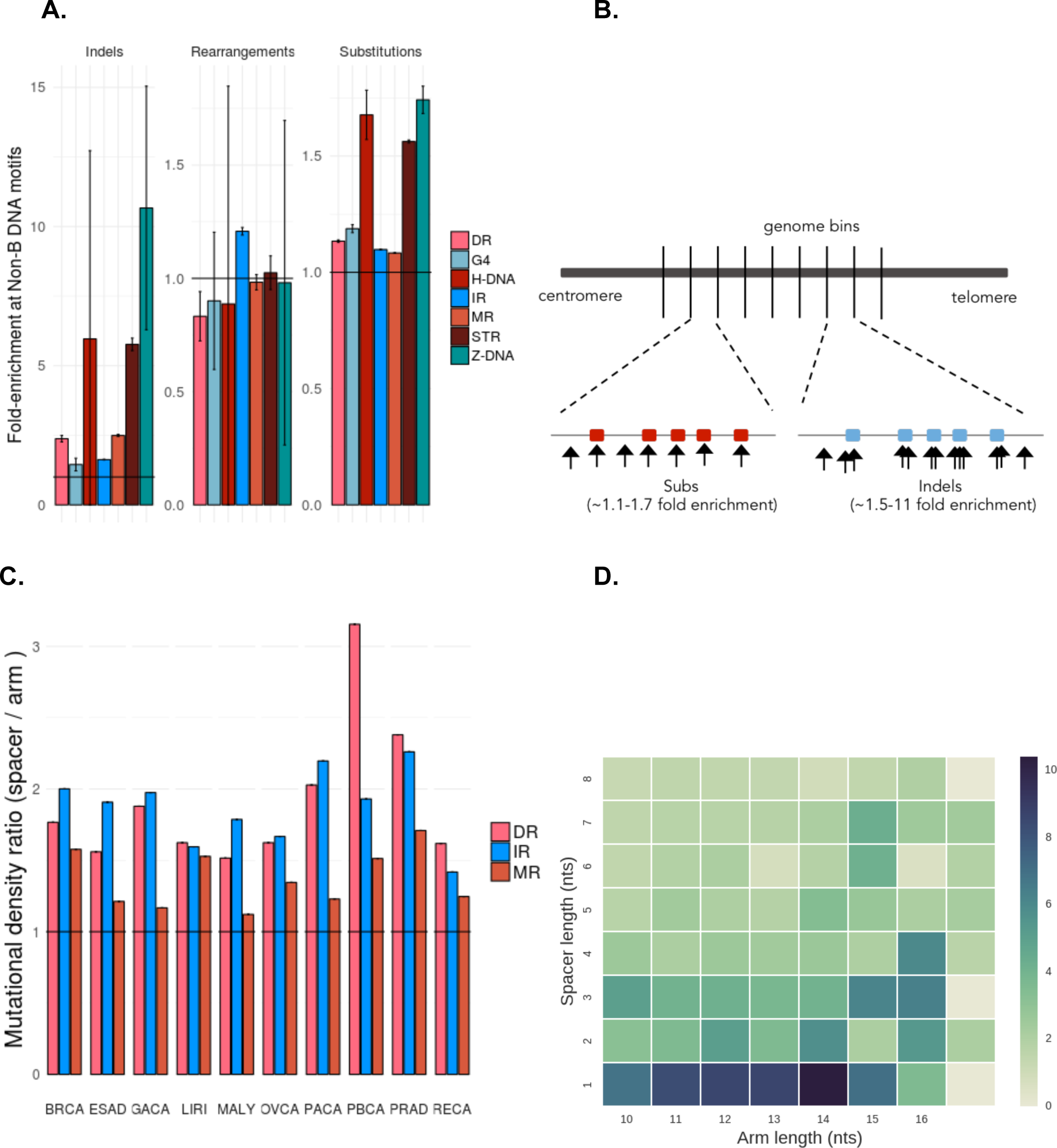
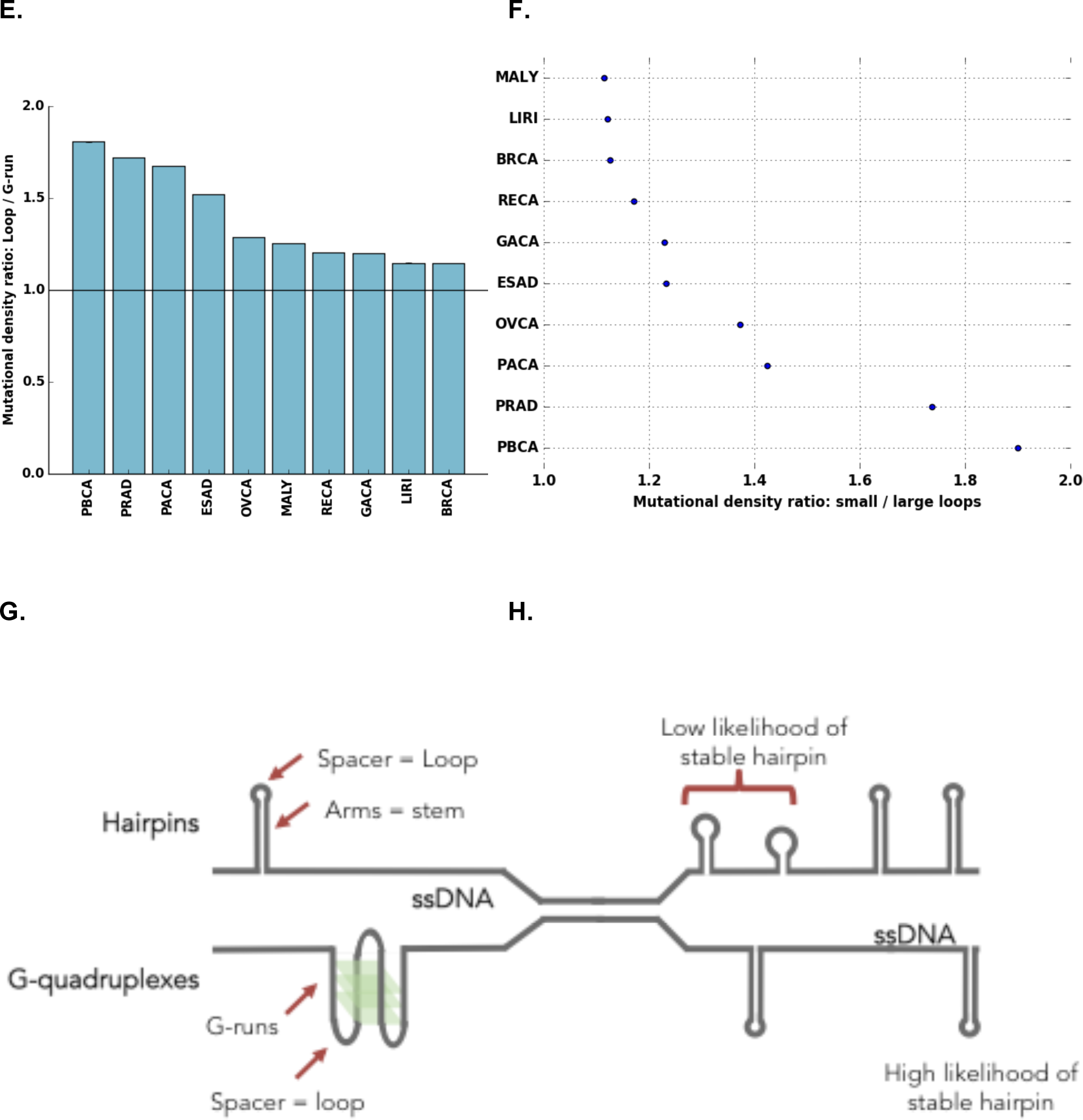
**Non-B DNA motifs are mechanistically linked to mutability through formation of secondary structures.** *(a)* Enrichment of mutagenesis for non-B motifs within their genomic bins, thus correcting for genomic GC variation. Error bars represent the standard error. *(b)* Depiction of enrichment per genomic bin, for results in (a), demonstrating how mutations are enriched for non-B motifs. Red and blue boxes represent non-B motifs. *(c)* Mutational density in spacers compared to arms for direct repeats, inverted repeats and mirror repeats across 10 tumour types. Error bars representing standard error are too small to visualise. Wilcoxon signed-rank test was performed (p-value <0.001 across all tumours for IR, MR, DR) *(d)* Heatmap of mutational density as a function of spacers and arms for breast cancer. *(e)* Enrichment of mutation density in loops: G-runs across ten cancer types. Error bars represent standard deviation from bootstrapping with replacement (n=10,000). *(f)* Enrichment of mutation density at G-quadruplexes for small loop sizes (less and equal to 3nt) relative to large loop sizes (more than 3nt) across ten cancer types. Error bars represent standard deviation from bootstrapping with replacement (n=10,000). Mann-Whitney U test was performed for each cancer type (p-value < 0.001 across all tumour types), (g) Depiction of two very different secondary structures that both have loop domains which are more mutable than their other components *(h)* Some non-B motifs have characteristics such as arm or spacer lengths that increase the likelihood of stable hairpin formation. There perhaps can occur stably more frequently and thus, their exposed regions are more likely to be damaged and mutated.

Second, we find that the elevated mutation densities in non-B motifs show domain-specificity. Selected non-B motifs have identifiable subcomponents - DR, IR and MR consist of two symmetric “arms” flanking a stretch of “spacer” sequence (Fig. 1 d-f). The arms can hybridize forming a transiently stable structure leaving the spacer sequence exposed to damage to potentially be more mutable (Fig. 1d-f). We find that spacer sequences are more enriched for substitutions than arm sequences (1.8-fold for DR, 2-fold for MR and 1.7-fold for IR) (Fig. 3c, Fig. S10, Fig. S11). This is in-keeping with previous experimental reports demonstrating how the loop domain formed by the spacer sequence tended to mutate more frequently in hairpin structures ^42,43^. It also reinforces a report that specifically explores a more conservative subset of IRs (with specific spacer and arm lengths), which suggests that mutability is an intrinsic property of these IRs, because nearly all mutational processes are elevated in IRs regardless of mutational process active in each tumor (Xueqing Zou et al., manuscript in submission).

Third, non-B motifs do not have a uniform thermodynamic capacity to form secondary structures. Experimental and biophysical simulation studies suggest that hairpin formation (for example) is optimal at certain spacer and arm lengths ^44,45,46^. If the physical formation of a secondary structure influenced mutability, then we would expect to observe elevated mutabilities particularly for spacer and arm lengths that are most favorable for hairpin/cruciform formation ^47,48^. Intriguingly, we find that spacer-to-arm mutation enrichment is indeed variable for different spacer sizes and various arm lengths (Fig. 3d, Fig. S10, Fig. S11). Heatmaps of mutation enrichment demonstrate that for IRs, which form hairpin and cruciform structures, mutability is greatest for spacer sequences of 1-3nt and arm lengths of 10-14nts (Fig. 3d) in-keeping with previous reports highlighting physical specifications of *in vitro* IR mutability ^47,48^. Also, DRs with short spacers and longer arms are more mutable (S10-S11) consistent with them being more likely to induce slipped structure misalignment^49^. By contrast, MRs exhibit more modest enrichment for particular spacer or arm lengths (Fig. S12). However, a small subset of MRs are H-DNAs that have high AG content (>90%) and are more likely to form triple-helical structures held together by Hoogsteen bonds (Fig. 1f). H-DNAs are believed to be more mutable than MRs^16^ and we do observe an excess of mutability in H-DNA in our analysis (Fig. 3a). These observations across IRs, DRs and MRs are recapitulated in other tumor types (Fig. S13).

Fourth, our findings are reinforced by assessing non-canonical secondary structures with very different physical properties. Primary sequence comprising G-runs and interspersed loop elements can form a complex G4 structure (Fig. 1c). Experiments in yeast systems have shown that smaller loop elements confer greater thermodynamic stability to G4 formation where the exposed loops are prone to mutation (Fig. 1c) ^50,51,52^. Indeed, our analysis supports these experiments showing that loops have a ~1.15-1.8 fold enrichment in mutagenesis over G-runs (Fig. 3e) and the subset of G-quadruplexes with average loop size of up to 3nt is more mutable than their counterparts with larger loop elements (Fig. 3f).

In conclusion, the relationship between somatic mutation and non-B motifs is not simply an association - we find incriminating evidence to suggest that it is the physical formation of secondary structures that predispose to damage and mutagenesis: Not only are non-B motifs enriched for mutation (Fig. 3a), the enrichment is domain-specific for selected non-B motifs (Fig. 3g), and biophysical characteristics that predispose to stable secondary structure formation (such as loop size and stem length) appear to be associated with increased mutability (Fig. 3h).

Non-canonical configurations are therefore primary determinants of mutagenesis - potentially raising the prior probability of mutability to considerable levels in a highly localised way, at specific locations. This has significant consequences for the biological interpretation of recurrent mutations.

A central tenet in cancer biology is the identification of driver mutations - those causally implicated mutations that are believed to drive tumorigenesis. Most drivers are found in protein coding sequences, although recent WGS studies permit the exploration of non-coding sequences^13,6,53,54^. Due to the difficulties of interpreting non-protein coding sequences, a useful criterion for identifying putative non-coding driver mutations is to focus on recurrently mutated loci^13,6^. We have demonstrated that non B-DNA motifs confer a marked propensity for increased mutability at local levels. Thus, we hypothesize that these motifs could be overrepresented amongst recurrently mutated loci. Indeed, one example of a statistically significant recurrently mutated locus is the promoter of the *PLEKHS1* gene that has been shown to be an inverted repeat^13,6^. For the cancer-types in our study, we first find that there are more recurrent substitutions than expected based on a truncated Poisson null model (Fig. 4a). Second, non-B DNA motifs are indeed overrepresented (five-fold) amongst recurrent substitutions (same site mutated two or more times) than non-recurrent ones (Fig. 4b-c, Fig. S14). Enrichment is variable from one motif to another, with short tandem repeats having 20-fold enrichment. Our finding that non-B DNA motifs are enriched for mutations, and in particular recurrent mutations, due to the formation of secondary structures (Fig. 4d) has important implications - effectively obfuscating the interpretation of recurrently mutated loci. Consequently, the cautionary note is this: statistical models of background mutability should consider the contribution to localised mutability provided by non-B DNA motifs in all future analyses.

**Figure 4:**
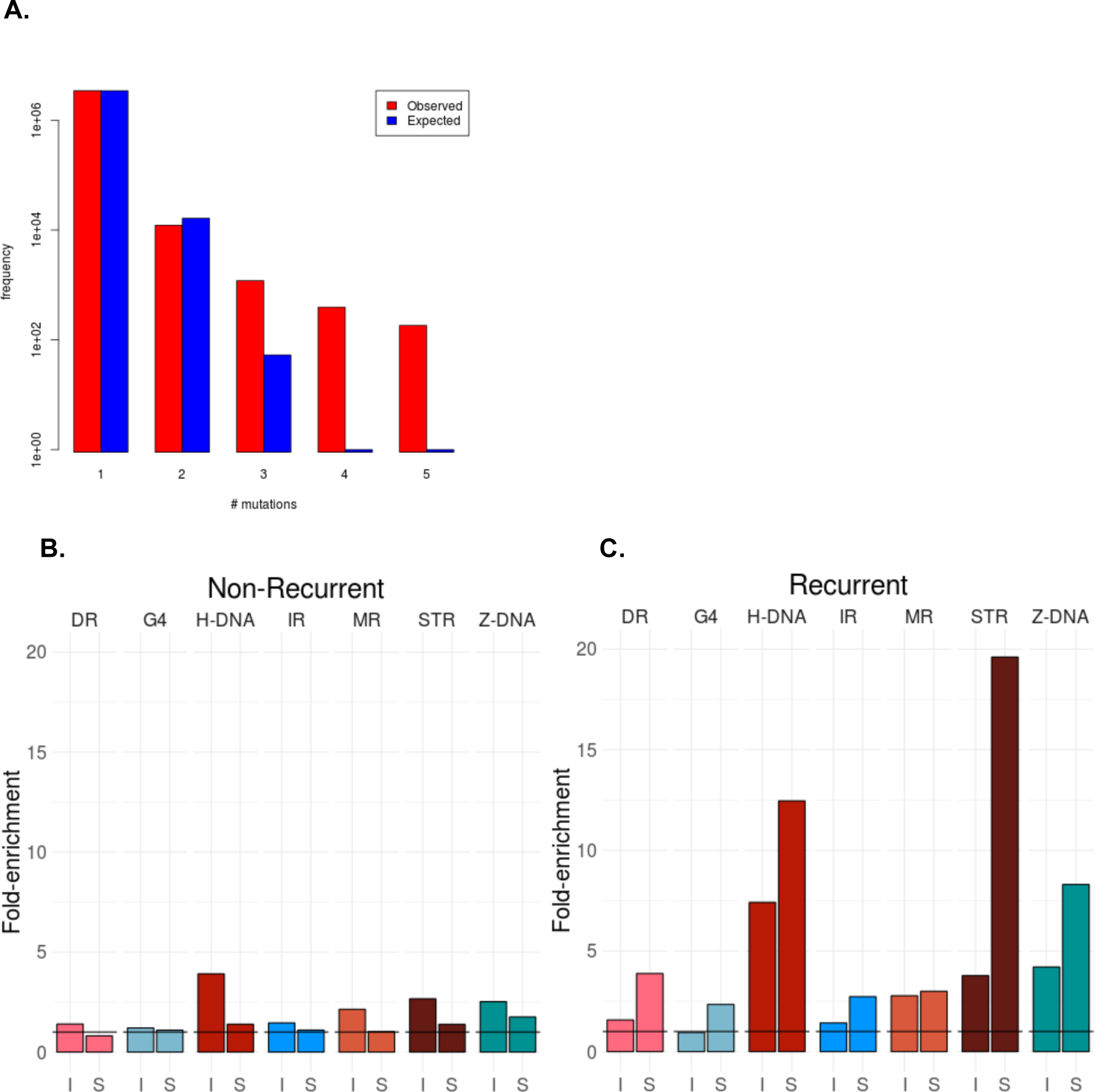
**Non-B motifs contribute to locally elevated mutation rates resulting in recurrent mutations the human genome.** (a) Distribution of the number of recurrent events for 3,476,890 somatic mutations from 560 breast cancers^6^. The values do not fit a truncated Poisson distribution (Chi2-test, p<1e-16) as there are more recurrent mutations than predicted by the null model. (b) Enrichment of non-recurrent mutations overlapping non-B-DNA motifs for indels (I) and substitutions (S). *(c)* Enrichment of recurrent mutations overlapping non-B DNA motifs for indels (I) and substitutions (S). Mann-Whitney U test for substitutions: p-value <0.001 for all non-B DNA motifs. Mann-Whitney U test for indels: p-value < 0.001 for STR, H-DNA, Z-DNA, MR and p-value <0.05 for DR and G4.

## Acknowledgements

This work has been performed on data that were previously published under the auspices of the International Cancer Genome Consortium and Breast Cancer Somatic Genetics Study (BASIS), a research project funded by the European Community's Seventh Framework Programme (FP7/2010-2014) under the grant agreement number 242006.

MH is supported by the Wellcome Trust Sanger Institute core grant. SN-Z was a Wellcome-Beit Fellow and personally funded by a Wellcome Trust Intermediate Clinical Research Grant (WT100183MA) at the start of writing this manuscript, and subsequently funded by a CRUK Advanced Clinician Scientist Award (C60100/A23916).

## Author contributions

IGS, MH and SNZ conceived the concepts and analytical framework, drove the intellectual exercise and wrote the manuscript. IGS, SM, NJ and MH wrote the code for analyzing and presenting the data.

## Materials and Methods

### 1. Somatic variants from cancer data

Data were obtained from whole genome sequenced cancers for breast cancer (n=560) from ^6^ and from 9 cancer types publicly available in ICGC^8^. The ICGC project codes for the cancer types were: PACA-CA (n=148)^55^ and PACA-AU (n=94) for pancreatic cancer^32^, OV-AU (n=72) for ovarian cancer^31^, LIRI-JP (n=264) for liver cancer^56^, PRAD-CA (n=120) for prostate cancer^33^, ESAD-UK (n=98) for esophageal adenocarcinoma^7^, GACA-CN (n=40) for gastric cancer^7^, RECA-EU (n=74) for renal cell cancer^8^, PBCA-DE (n=239) for pediatric brain cancer^8^ and MALY-DE (n=100) for malignant lymphoma^8^. In total, 1809 whole genome sequenced cancers were analysed. Sequencing coverage exceeded 25X for all tumours and matched normal samples.

Short insert paired-end reads were aligned to the reference human genome (GRCh37) using Burrows-Wheeler Aligner, BWA (v0.5.9).

High quality curated somatic variant calls (substitutions, insertions/deletions and structural variations) were derived from the Wellcome Trust Sanger Institute's Cancer Genome Project whole genome sequencing pipeline as previously described^6^. This is constituted by a bespoke, Expectation-Maximisation- based substitution-calling algorithm (CaVEMan)^57,58^, a modified version of an insertion/deletion detection algorithm, Pindel^59^ and a bespoke structural variant algorithm which uses de Bruijn graphing for discovery of somatic rearrangements and local reassembly for mapping breakpoints to base pair level.

A subset of all somatic variants for breast cancer samples had been previously validated using alternative sequencing platforms to ensure high specificity of data^6^. In short, 70 samples were used for validation across a mix of histopathological subtypes and were sourced from different collaborating centres.

- On average 3% (range 0.6-20%) of the total burden of substitutions per sample were used for validation (total 11,581 mutations). The positive predictive value was ~95.5% (average) for substitutions.
- On average of 40% (range 8%-68%) of the total number of indels were validated per sample (total 7,192). The positive predictive value was 85% for indels.
- Rearrangements were discovered using Brass I and an additional *in silico* method was used *(de novo* breakpoint assembly) to validate the finding. Only breakpoints that were *de novo* assembled with high confidence (80% and above only) were included in order to reduce the likelihood of false positive calls. PCR-based Sanger sequencing validation confirmed the presence of 803 randomly sampled breakpoints from this conservative dataset.

**Table 1:** **Number of substitutions, indels and rearrangement breakpoints per tumour type.**

| Cancer Name | Samples | Substitutions | Indels | Rearrangement breakpoints |
| --- | --- | --- | --- | --- |
| BRCA | 560 | 3,479,651 | 371,993 | 131,068 |
| LIRI | 264 | 3,575,056 | 852,361 | 51,034 |
| OVCA | 72 | 732,189 | 141,296 | 39,078 |
| ESAD | 98 | 2,890,654 | 347,680 | 48,394 |
| GACA | 40 | 525,850 | 185,213 | 12,268 |
| PBCA | 239 | 299,241 | 231,874 | 13,120 |
| PACA | 242 | 1,881,336 | 625,803 | 48,404 |
| RECA | 74 | 584,144 | 123,180 | 1,972 |
| MALY | 100 | 1,242,356 | 203,051 | 10,752 |
| PRAD | 120 | 602,729 | 799,583 | 24,104 |

Furthermore, these datasets have all been published and therefore been through peer-review previously.

### 2. Reference non-B DNA annotations

Non-B DNA sequence motif annotations were derived from^11^. We have focused on the following categories in this analysis: Mirror repeats, H-DNA, short tandem repeats, Z-DNA, inverted repeats, direct repeats and G-quadruplexes.

- A mirror repeat is a section of sequence that is repeated with a center of symmetry on the same strand, length of at least 20nt and arm size of at least 10nt. A subset of mirror repeats are termed Hinged DNA (H-DNA), because they are predisposed to forming a triple helical structure connected through alternative chemical bonds called Hoogsteen bonds. H-DNA have a high (>90%) AG content, arm lengths of >=10nt and spacer size of less than 8nt.
- Z-DNA is a left-handed double helical structure that is formed by alternating purine-pyrimidine tracts of at least 12nt (excluding AT repeats).
- Direct repeats are defined as repeated sequences with arm length of >=10nt, with maximum size 300nt.
- Short tandem repeats are defined as motifs of 1-9nt, repeated at least 3 times with a minimum length of 9nt and without any interruptions. Short tandem repeats are prone to misalignment and formation of looped or slipped structures.
- Inverted repeats are palindromic sequences with minimum arm length of 6nt, spacer size up to 100nt and have a tendency to form hairpin or cruciform structures.
- G-quadruplexes are defined as 4 or more runs of at least 3 guanines, separated by spacers of 1-7nt of other nucleotides. For G-quadruplex motifs we referred to G-runs as the guanine runs that can form Hoogsteen bonds and the loops as the spacer between the G-runs.

Bedtools utilities v2.21.0 were used to manipulate genomic files and intervals.

To count the number of nucleotides shared between different non-B DNA motifs bedtools intersect and coverage functions were used. Bedtools jaccard was used to calculate the Jaccard index for each pair of motifs (Fig. S1b).

### 3. Epigenomic Data

DNase and histone modification narrowpeak files were downloaded from Roadmap Epigenomics Mapping Consortium data coordination center (<u>http://www.roadmapepigenomics.org/data/</u>) and BAM files were derived from ENCODE Project repository (<u>http://genome.ucsc.edu/ENCODE/downloads.html</u>). HMEC cell line epigenetic narrowpeak data were used to model breast cancer, whereas PANC1, HepG2 and GM12878 cell line narrowpeak epigenomic data were used to model pancreatic cancer, liver cancer and malignant lymphoma. Ovary, esophagus, fetal female brain, stomach mucosa, fetal kidney primary tissue narrowpeak epigenomic data were used to model ovarian cancer, esophageal carcinoma, pediatric brain cancer, gastric cancer and renal cell cancer respectively. BAM files for the same epigenetic modifications were derived from ENCODE consortium to validate the findings derived from narrowpeak files, for MCF-7 cell line which is used to model breast cancer.

### 4. Chromatin States

Chromatin states represent partitions of the genome derived using epigenetic data. Here they were defined with Segway as described in ^60,61^.

The background or “expected” density of each non-B DNA motif (DN-background) was calculated as the total number of occurrences of the motif (TO) over all mappable nucleotides across the Segway states (TN). The density of each non-B DNA motif at a particular state (DN-specific) was calculated as the fraction of the number of occurrences of the non-B DNA motif at that state (SO) over the number of mappable nucleotides covered in that state (SN). The enrichment of a non-B DNA motif at a given state was the fraction of the density at the state over the background density of the motif.

Background density of a non-B DNA motif: DN-background: TO /TN

Density of a non-B DNA motif at the state = DN-specific: SO /SN

Enrichment: DN-specific / DN-background

The mean enrichment across 6 human cell lines (GM12878, H1-Hesc, HepG2, HUVEC, K562, HelaS3) was calculated. Hierarchical clustering of chromatin states and plotting was performed with the python package “Seaborn” using default parameters (Fig. 1i).

### 5. Repli-Seq Data

Reference coordinates for replication landmarks were inferred from Repli-Seq data of 14 cell-lines, which were NHEK, IMR90, HUVEC, HeLa-S3, GM12813, GM12812, GM12801, GM06990, BJ, BG02ES, MCF-7, GM12878, HepG2 and K562. Repli-Seq data were obtained from the ENCODE project (<u>https://www.encodeproject.org/</u>) and processed as described in^4^. Replication timing was measured at each genomic interval using bedtools map utility function. Repli-Seq data for MCF-7 were used for breast cancer, HepG2 Repli-seq data were used for liver cancer, GM12878 for malignant lymphoma and MCF-7 for all other cancer types. A positive correlation between mutations and replication time indicates positive correlation for early replication time domains and mutations, while a negative correlation denotes a positive correlation for late replication time domains (Fig. 2a, Fig. S4). Pearson correlation between any two cell lines with Repli-Seq data exceeded 0.69 in all cases, using 500kB genomic windows (Fig. S2).

### 6. Modelling the relationship between mutations and genomic features (epigenetics, replication time domains and non-B DNA motifs)

The human genome (hg19) was partitioned in equal-sized regions of 500kB segments. Centromeric sites, simple repeats and regions of excessive sequencing depth (UCSC Top 0.01 Hi Seq Depth) were downloaded from the UCSC genome browser and used to identify bins with low mappability. Bedtools coverage was used to calculate the coverage of centromeric sites and low complexity sequences at each genomic interval. We excluded the first and last bin from each chromosome as well as any bin where <50% of the bases were mappable or where replication time data is missing, and the sex chromosomes. This resulted in 5,581 non-overlapping bins. All quantities except for the replication times were transformed as x' = log2(1 + x) for the downstream analysis.

To map reads from histone modification BAM files from ENCODE consortium at each genomic interval, bedtools multicov utility function was used. In case of multiple replicates per file, the mean number of reads per segment was calculated across replicates.

To calculate the number of non-B DNA motifs, mutations, genomic features and narrowpeak files from Roadmap epigenetic consortium at each genomic segment, bedtools intersect utility was used (bedtools intersect −a segments.file −b mutation.file with flags −u, −v, −c).

Bedtools nuc algorithm was used to calculate the GC content at each interval as well as the number of As, Gs, Cs, Ts and Ns at each interval for the hg19 reference genome.

Partial correlation is a measure of association between two variables, controlling for the effect of covariates. Partial correlations were applied to measure the relationship between mutations and non-B DNA motifs, controlling for the effect of epigenetic markers and replication timing. Partial correlations were calculated in R with the package 'ppcor'. Results are noted in Fig. S3.

### 7. Linear and random forest regression

To model the relationship between the number of mutations and a plethora of explanatory variables we applied two predictive models; linear regression and random forest regression. In the former, additive relationships are modeled using linear predictor functions, whereas a random forest model is an ensemble learning model in which multiple regression trees are constructed and evaluated. In both models, the relative importance of each predictor variable can be measured. The two models were applied independently to each cancer type. Prostate cancer, for which epigenetic data from a relevant cell of origin were not available, was excluded.

Both models were evaluated using 10-fold cross-validation, whereby the model was trained using 90% of the data and tested using the held out 10%. The same bins and transformations that were used for the correlation calculations were used for the regression. For the linear model we used the command “lm” in R and for the random forest regression, we used the R-package “randomForest” with default parameters. For the random forest regression model feature importance was measured using the predictive measure of the original and the permuted dataset. In particular, the variable importance for Fig. 2 panels c and d was evaluated using the R-package “pRF", which uses a permutation test. The parameters for the pRF function were “n.perms = 200” and “mtry = 4". The biplot in Fig. 2e represents the first two loadings obtained using the princomp command in R.

### 8. Enrichment of mutagenesis within non-B DNA motifs

For each bin of size *B* and non B-DNA motif, we calculated the number of bps covered, *b,* as well as the number of mutations that overlapped the motif type, *m,* and the number of mutations not overlapping the motif, *n.* The fraction of mutations overlapping non B-DNA motifs is *m/b,* and the fraction of mutations not overlapping motifs is *n/(B-b).* The enrichment of mutations overlapping non B-DNA motifs is given by *r = m(B-b)/nb.* When calculating *r*we exclude the bins where *b = 0*. When calculating ratios in Fig. 3a, Fig. S9, the expected values and the variances are adjusted to account for correlations as:

> E[X/Y] = E[X]/E[Y] − Cov[X, Y]/E[Y]^2^ + Var[Y]E[X]/E[Y]^3^

and

> Var[X/Y] = (E[X]/E[Y])(Var[X]/E[X]^2^ − 2Cov[X,Y]/E[X]E[Y] + Var[Y]/E[Y]^2^).

For panels in Fig. S10, Fig. S11 this correction was not applied since it results in negative values for some of the spacer or arm lengths. For all cases, the correlation between the mutation densities of spacer and arms is positively correlated and for all but a handful, the adjusted ratios are an order of magnitude higher than the unadjusted, suggesting that the latter is an underestimate.

When discussing mirror repeats, direct repeats and inverted repeats “spacer” is used to denote the part of the motif that is not repeated, whereas “arm” is used to denote the repeating parts (Fig. 1 d-f). The number of mutations overlapping spacers and arms was recorded separately.

For each direct repeat, inverted repeat and mirror repeat motif we calculated the length of the spacer and the arms. The mutation density was calculated as the number of mutations overlapping each motif part divided by the length of the spacer or arm respectively, averaged across all instances of the motif-type. Figure 3c is a summary figure across the ten tumour types of Fig. S10 and Fig. S11, measuring the average mutational density in the spacer and arm for spacer sizes 1-10nt.

We measured the mutational density in each sub-component of the G-quadruplex motif (G-run and loop) and calculated the enrichment as the fraction of the mutational densities of the two sub-components. Furthermore, we separated G-quadruplexes into two groups based on the average size of the loops (less or equal than 3nt or longer than 3nt) and compared the mutational density of each group. In all cases, error bars displayed standard deviation measured using bootstrapping with replacement (n=10,000), with a custom python script.

### 9. Analysis of recurrent mutations

The number of somatic mutations, indels and rearrangements at the each genomic site was calculated per cancer type across patients using a python script, which is available upon request. The overlap between recurrently mutated sites for each mutation type and each non-B DNA motif was subsequently calculated using bedtools intersect utility. A truncated poisson model was applied as the null model.

The truncated Poisson model was estimated using the “mle” function from the “stats4” R-package. Mann-Whitney test for recurrent and non-recurrent indels and substitutions overlapping each non-B DNA motif was calculated to measure significance.

